# Multiplexed bioluminescence microscopy via phasor analysis

**DOI:** 10.1101/2021.06.18.448905

**Authors:** Zi Yao, Caroline K. Brennan, Lorenzo Scipioni, Hongtao Chen, Kevin Ng, Michelle A. Digman, Jennifer A. Prescher

## Abstract

Microscopic bioluminescence imaging has been historically challenging due to a lack of detection methods and easily resolved probes. Here we combine bioluminescence with phasor analysis, an optical method commonly used to distinguish spectrally similar fluorophores. Bioluminescent phasor enabled rapid differentiation of multiple luciferase reporters and resonance energy transfer processes. The merger of bioluminescence and phasor analysis provides a platform for routine, time-lapse imaging of collections of cellular features.

## MAIN

Bioluminescence imaging (BLI) with luciferase enzymes and luciferin small molecules has been used for decades to interrogate biological processes in vivo.^1^ Unlike fluorescence, BLI requires no excitation light and can thus be advantageous for visualizing processes in tissues and whole organisms.^2^ Bioluminescent probes can be imaged repeatedly over time without detriment, as there are no phototoxicity concerns. Brighter and more tissue penetrant luciferase-luciferin pairs are also continuing to push the boundaries of what can be “seen” in large organisms.

While ubiquitous for whole animal studies, BLI is rarely used for imaging at the microscopic scale. This is due, in part, to a lack of suitable reporters and detection methods. Unlike fluorescence imaging, where large collections of probes and multi-parameter imaging methods exist,^3^ BLI has fewer reporters that can be used concurrently.^1^ The lack of well-resolved probes precludes routine multiplexed imaging. BLI has also been stymied by weak photon outputs from conventional luciferases. Dim reporters require long acquisition times, resulting in hazy images that fail to capture dynamic events.^4^

The BLI landscape has changed dramatically in recent years with the development of Nano-lanterns,^5^ bioluminescent reporters that are orders of magnitude brighter than conventional probes and detectable on standard microscopes.^6-7^ BLI with Nano-lanterns rivals the spatiotemporal resolution of fluorescence microscopy,^7-8^ but a major hurdle remains: differentiating the probes. Nano-lanterns, like other luciferases, exhibit broad emission spectra, complicating their resolution by color alone. Further resolution can be achieved using luciferases that respond to distinct luciferin scaffolds, but longer imaging times are required as the substrates cannot be used simultaneously. Signal from the first probe must dissipate before the second luciferin is added.^9^

Here we address the need for more effective multiplexing at the microscale by developing bioluminescent phasor—a technology that combines the exquisite sensitivity and dynamic range of BLI with the resolving power of phasor imaging. Phasor analysis can deconvolute highly overlapping spectra (common among bioluminescent reporters).^10-11^ The method is superior to conventional spectral unmixing algorithms for resolving weak emitters owing to minimized spectral noise in phasor transform.^12^ To establish bioluminescent phasor imaging, we built a camera-based microscope using widely accessible components. Six reporters were easily resolved in a single live-cell imaging experiment—a record in bioluminescence imaging. The readouts were quantitative and instantaneous, enabling bioluminescent components to be readily assigned in a given image. Furthermore, unlike conventional spectral^13^ and substrate unmixing,^9^ no *a priori* information on the reporters themselves was necessary for deconvolution. Bioluminescent phasor also takes advantage of the full spectrum of luciferase emission, enabling facile assignment of engineered probes and even BRET efficiency.

Since bioluminescent photons derive from distinct luciferase-luciferin interactions, we surmised that each pair would produce a unique phasor output, dictated by specific enzyme-substrate interfaces (Fig. 1a). Unique wavelengths and spectral shapes would register as distinct signals on a 2D plot, while mixtures would register as linear combinations of the clusters. The approach is analogous to delineating collections of fluorescent emitters using phasor analysis.^11^ The workflow for measuring bioluminescent signals featured an all optical, four-channel acquisition scheme based on a previously described sine and cosine filter setup (Fig. 1b and Supplementary Fig.1).^14^ Briefly, the emission light was split into four channels, with two of the channels passing through sine and cosine filters. Intensities collected from the sine and cosine channels were computationally referenced with non-filtered channels to provide bioluminescent phasors with high resolution. All necessary spectral information for computing phasor locations was captured in a single frame (in 10 s or less). This feature is crucial as bioluminescent probes exhibit fast-decay signal dynamics that complicate sequential scanning over multiple windows.^4^ Importantly, the acquisition scheme can be easily implemented with standard camera-based detection platforms (e.g. microscopes), as shown in this study.

**Figure 1.**
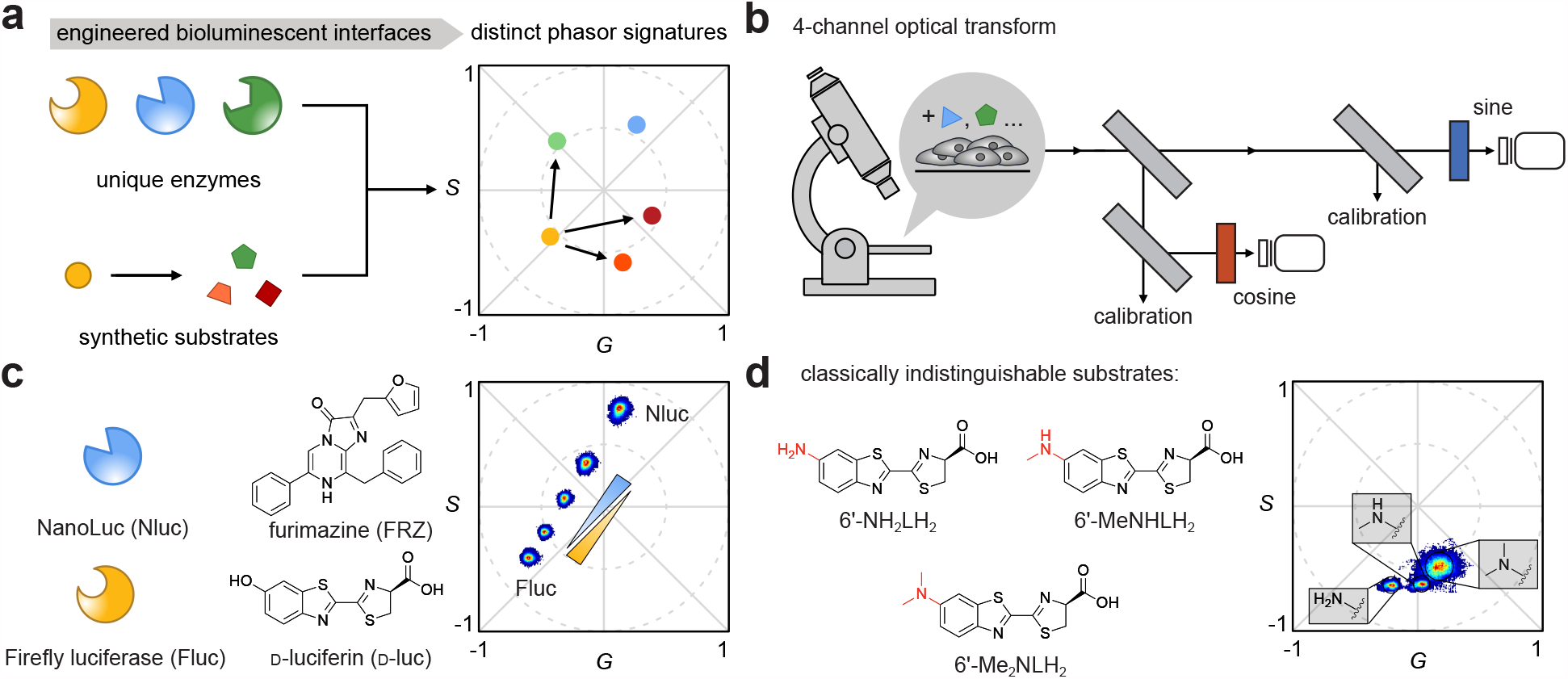
Bioluminescent phasor imaging. (a) Distinct luciferase-luciferin pairs produce unique phasor signatures. (b) A 4-channel detection scheme for capturing bioluminesence. (c) Common luciferase reporters and (d) structurally similar luciferin analogs produced distinguishable fingerprints on the phasor plot.

Using the 4-channel setup, we first evaluated the bioluminescent phasors of different luciferase-luciferin pairs. Firefly luciferase (Fluc) and its complementary luciferin (D-luc) were readily resolved from NanoLuc (Nluc) and its substrate (FRZ). Solutions of the two bioluminescent probes generated unique clusters on the phasor plot that matched the expected emission properties (Supplementary Fig. 2). Different ratios of Fluc and Nluc registered as clusters along a line between the two pure populations (Fig. 1c and Supplementary Fig. 2). Notably, both luciferins were added in a single bolus. Simultaneous BLI is often complicated due to spectral overlap and enzymatic crosstalk among the probes.^1,9^ Multi-component imaging can be achieved adding one luciferin at a time, but such processes typically require multiple hours to complete. Signal from one luciferin must diminish before the second is administered, precluding dynamic studies of gene expression and other features. Phasor imaging, by contrast, enables simultaneous and continuous visualization. Phasor locations for other well-characterized luciferase-luciferin pairs^15^ were also readily differentiated (Supplementary Fig. 3).

**Figure 2.**
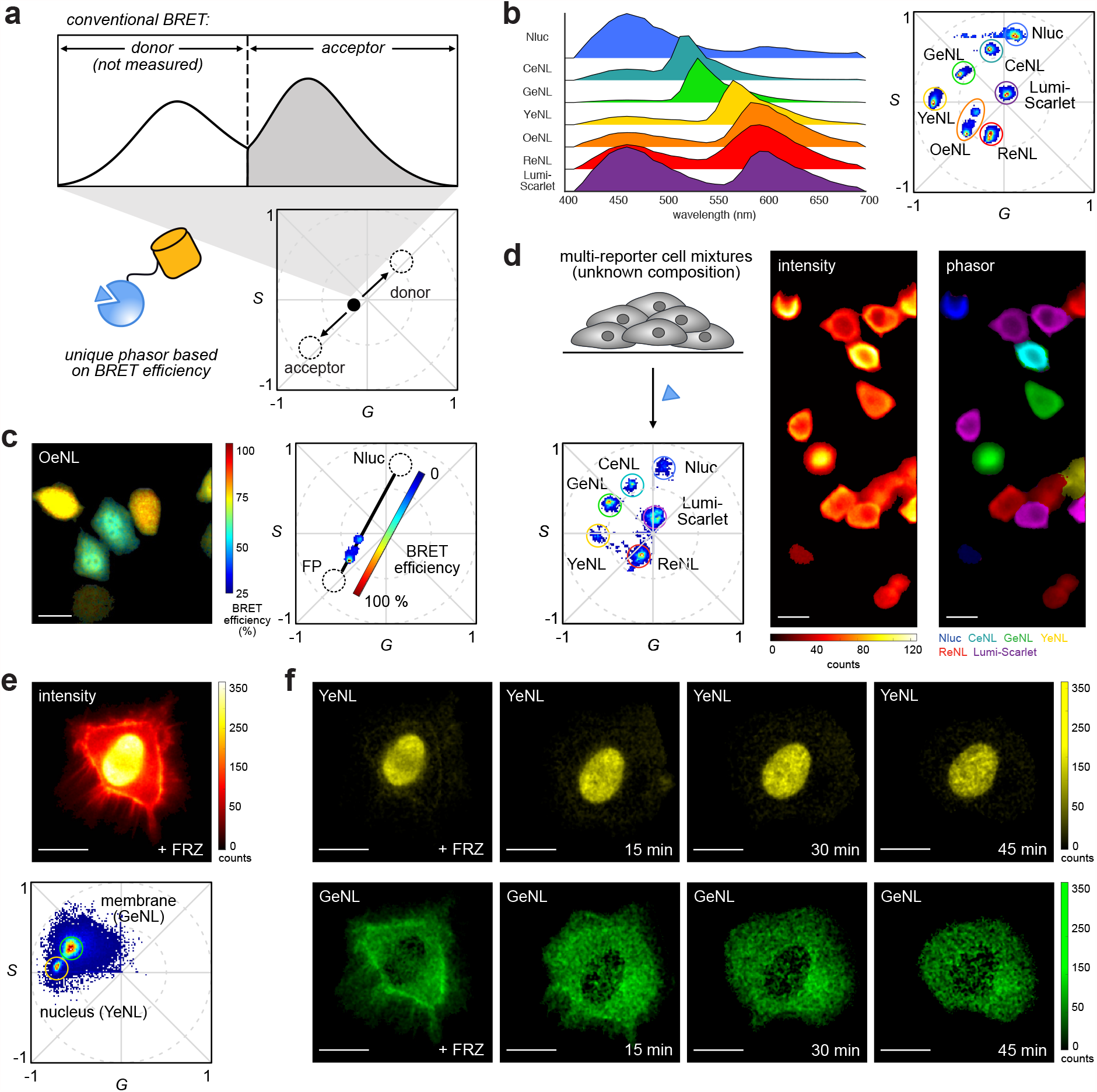
Facile multicomponent imaging with BRET probes and bioluminesent phasor. (a) Unique phasor signatures can be generated from incomplete energy transfer. (b) A collection of seven BRET reporters were readily distinguished on the phasor plot. (c) Phasor signature of OeNL revealed heterogeneous BRET efficiencies with single cell resolution. (d) Cell mixtures were identified using the phasor fingerprints of individual reporters. (e) Simultaneous tracking of two subcellular features using unmixed phasor signals. For (c)-(f), the scale bar represents 25 μm.

Bioluminescent phasor could resolve unique enzyme-substrate interfaces resulting from minimally modified luciferin analogs. D-Luc analogs comprising small steric appendages^16^ and/or amino substituents^17^ are attractive imaging agents owing to their robust and often red-shifted output. However, these probes are classically inseparable due to their overlapping emission (λ_max_ = 594-623 nm) and broad spectra (80-90 nm bandwidth at half maximum).^17^ The analogs exhibit a range of binding interactions with Fluc, though, that modulate the spectral output and register as unique phasor signatures. Indeed, distinct signals were observed when the analogs were combined with Fluc (Fig. 1d). Even regioisomers with similar light-emitting cores could be distinguished, highlighting the resolving power of the approach and its capacity for multiplexing^18^ (Supplementary Figs. 4-6).

We further examined whether phasor analysis could be interfaced with genetically encoded BRET reporters. These agents comprise a luciferase donor and fluorescent protein acceptor that emit light in a distance-dependent fashion. BRET reporters are widely used,^4^ but conventional applications rely on filters to capture the acceptor emission, resulting in signal loss and often poor resolution. Multiplexing probes with low BRET ratios is also challenging, as the broad, residual emission from the donors can overwhelm outputs from the acceptors, precluding signal differentiation. We hypothesized that bioluminescent phasor could take such “undesired” spectral features into account, registering incomplete energy transfer as part of the reporter signature (Fig. 2a). Moreover, for a given luciferase-fluorescent protein (FP) combination, the position of the phasor between the donor and acceptor signals should report on BRET efficiency. Identical probes exhibiting different BRET ratios would be geographically distinct on the phasor plot. Such readouts on energy transfer efficiency could provide critical insights into single cell dynamics, among other processes.

To showcase the potential of bioluminescent phasor for BRET imaging, we transfected HeLa cells with a collection of common BRET constructs. Unique phasor signals were observed upon FRZ treatment, consistent with the expected emission spectra (Fig. 2b and Supplementary Fig. 7). Even probes with similar acceptor fluorophores (e.g., ReNL, OeNL and LumiScarlet, Supplementary Fig. 8) could be discerned based on their unique phasor signatures.^19^ For red-emitting BRET fusions (e.g., OeNL and ReNL) different cells produced distinct phasor clusters (Fig. 2c and Supplementary Fig. 9), likely due to lower donor-acceptor spectral overlap.^20^ The cell-to-cell heterogeneity likely reflects differences in BRET efficiency across individual cellular environments. The differences register as unique phasor signatures, highlighting the sensitivity of the approach in resolving BRET emission profiles. Signal assignments based on energy transfer efficiency could provide critical insights into single cell dynamics.

Bioluminescent phasor can readily resolve mixtures of cell populations and subcellular features. HeLa cells expressing unique BRET fusions were mixed and treated with FRZ. Multiple images were recorded, and each pixel was pseudo-colored according to the distinct phasor cluster. Cell populations with 3-6 unique reporters were readily distinguished (Fig. 2d and Supplementary Fig. 10). The identities of the encoded cells were determined by simple referencing to the phasor locations of pure populations, without any prior knowledge of the cell mixture composition. Additionally, probe combinations not easily differentiated by emission maxima alone, including GeNL/YeNL (λ_max_ = 539, 548 nm) and ReNL/LumiScarlet (λ_max_ = 602, 610 nm), were readily discerned (Supplementary Fig. 10). Subcellular compartments were visualized with high spatial resolution using localized YeNL reporters^13^ (Fig. 2e and Supplementary Fig. 11). The subcellular features were further enhanced via blind deconvolution and computational removal of out-of-focus light. Two-component imaging was performed by tagging the plasma membrane and nuclei of HeLa cells with GeNL and YeNL, respectively. The dynamic motions of the organelles were also continuously tracked for one hour (Fig. 2f and Supplementary Video 1). This experiment marks the first demonstration of simultaneous bioluminescent tracking of two subcellular features with no time delay.

In summary, we developed a new platform for multicomponent microscopy that leverages bioluminescent probes and phasor analysis. A camera-based system was repurposed to measure light emitted from luciferase-luciferin reactions, and phasor locations were determined computationally. Multi-component imaging was achieved with luciferases in solution and in live cells. Subcellular structures and dynamics were readily visualized. The workflow and instrumentation used to obtain bioluminescent phasors can be integrated with commercial microscopes and imagers. Time-lapse imaging with this excitation-free platform will enable many studies of dynamic events (e.g. optogenetics and circadian rhythms) that cannot be accessed via conventional fluorescence imaging. Bioluminescent phasor employs widely available optical components and user-friendly probes, making the technology easily accessible to non-specialists. We anticipate that the technique will provide new insights into complex biological networks and spur new directions in probe development.

## Acknowledgments

This work was supported by the U.S. National Institutes of Health (R01 GM107630 to J.A.P., P41-GM103540 to L.S. and M.A.D.) and the Paul G. Allen Frontiers Group (to J.A.P and M.A.D). Z.Y. was supported by the National Science Foundation via the BEST IGERT (DGE-1144901) program and a Graduate Research Fellowship (DGE-1321846). C.K.B was supported by the UCI Physical Sciences Machine Learning NEXUS program. We thank Prof. Enrico Gratton and members of the Laboratory of Fluorescence Dynamics (LFD, UCI) for helpful discussion. Some experiments were performed at the Laser Spectroscopy labs (LSL) at UCI.

## Author contributions

Z.Y., C.K.B., M.AD., and J.A.P. conceived the project idea. Z.Y., C.K.B., L.S., H.C. performed the experiments. L.S. and H.C. designed, and built the imaging setup. Z.Y. and C.K.B. prepared the biological samples. L.S. wrote the codes and analyzed the data. K.N. generated the reporter constructs. All authors analyzed data and contributed to the writing of the manuscript. All authors have given approval to the final version of the manuscript.

