## Supporting Information for "Multiplexed bioluminescence microscopy via phasor analysis"

### MATERIALS AND METHODS

#### Reagents

All reagents were purchased from commercial suppliers were of analytical grade used without further purifications. 4′-BrLuc, 5′-BrLuc, 7′-BrLuc, 4′-MeLuc, 7′-MeLuc, 6′-$NMe_2$Luc, 6′-NHMeLuc, 6′-$NH_2$Luc, 5′-Pyr-Luc and 7′-Pyr-Luc were prepared as previously described.[16,17, 18]

#### Mammalian cell culture and transient transfection

HeLa cells (ATCC) were cultured in complete media: DMEM (Corning) containing 4.5 g/L glucose, 2 mM L-glutamine, and supplemented with 10% (v/v) fetal bovine serum (FBS, Life Technologies), penicillin (100 U/mL), and streptomycin (100 μg/mL, Gibco). Cells were incubated at 37 °C in a 5% $CO_2$ humidified chamber, and were serially passaged using trypsin (0.25% in HBSS, Gibco). Cells were counted using an automated cell counter (Countess II, Invitrogen)

Cells (2.5 x $10^5$) were seeded in 12-well dishes (Corning). Transfections were performed with luciferase constructs using Lipofectamine 2000 or 3000 (Thermo Fisher) according to the manufacturer's instructions. Transfections were performed when cells were 75-80% confluent (1 day post seeding). The following plasmids were used for transfection, and were gifted from T. Nagai: CeNL (Addgene plasmid no. 85199), YeNL (Addgene plasmid no. 85201), YeNL-actin (Addgene plasmid no. 89521), YeNL-ER (Addgene plasmid no. 89516), YeNL-mito (Addgene plasmid no. 89517), YeNL-H2B (Addgene plasmid no.89522), OeNL (Addgene plasmid no. 85202), and ReNL (Addgene plasmid no.85203). Antares was a gift from M. Lin (Addgene plasmid no. 74279). LumiScarlet was a gift from H.-w. Ai. GeNL, Nluc-IRES-GFP and CD8LS-YeNL-CD4 plasmids were cloned using the general cloning methods mentioned below. Each plasmid was transformed into TOP10 competent *E. coli* cells. The transformants were plated on agar plates containing carbenicillin (50 μg/mL). Colonies containing the genes of interest were expanded overnight in 5 mL LB broth containing ampicillin (50 μg/mL). Plasmid DNA was extracted using a Zymo Research Plasmid Mini-prep Kit and concentrations were measured using a Nanodrop 2000c Spectrophotometer (Thermo Scientific). Approximately 16 h post transfection, cells were lifted with trypsin and plated (5 x $10^5$) in tissue-culture treated 8-well chambered coverslips (μ-Slide 8 Well ibiTreat, Ibidi). The coverslips were coated with 5 mg/$cm^2$ fibronectin human plasma (Millipore Sigma) according to the manufacturer's instructions.

#### Luciferase expression and purification

Luciferases were expressed and purified as described by Rathbun, *et al.*[16] The YeNL-pCold, Fluc-pET28a, Nluc-pET28a, and Akaluc-pET28a plasmids were generated using the general cloning methods mentioned below. The Nano-lantern pRSETb plasmid (Addgene plasmid no.51969) was a gift from T. Nagai. Plasmids were transformed into chemically competent BL21 *E. coli* cells. The transformants were plated on agar plates containing kanamycin (Kan, 40 μg/mL). Cells were expanded in LB media containing Kan (LB-Kan) at 37 °C overnight. The overnight culture (20 mL) was used to inoculate 1 L LB-Kan. The new culture was incubated at 37 °C to mid-log phase (O.D. ~0.8). The culture was then induced with isopropyl β-D-1-thiogalactopyranoside (IPTG, 500 μM final concentration), and incubated at 22 °C for 16–18 h. Cells were harvested at 4 °C by centrifugation at 4000 rpm for 15 min. Cell pellets were resuspended in 40 mL of phosphate buffer (50 mM phosphate, 300 mM NaCl, 1 mM dithiothreitol (DTT), and 1 mM phenylmethylsulfonyl fluoride, pH = 7.4). Lysozyme (2 mg) was added, and the cells were sonicated and centrifuged at 10000 rpm for 1 h at 4 °C. Fluc, Nluc, Akaluc, YNL, and YeNL were purified from clarified supernatants using nickel affinity chromatography (BioLogic Duo Flow Chromatography System, Bio-Rad). Fluc and Akaluc were dialyzed into a Tris-acetate buffer (25 mM Tris-acetate, 1 mM EDTA, and 0.2 mM ammonium sulfate, pH = 7.8) at 4 °C for 16 h. Nluc, YNL, and YeNL were dialyzed into a phosphate buffer (50 mM phosphate, pH 7.8). DTT (1 mM final concentration) and

15% glycerol were added to the dialyzed samples prior to storage at –20°C. Final protein concentrations were determined using absorbance measurements at 280 nm collected on a JASCO V730 UV-vis spectrophotometer. SDS-PAGE analyses were also performed to verify protein purify. Gels were stained with Coomassie R-250.

**General cloning method.** Polymerase chain reaction (PCR) methods were performed to isolate and amplify all genetic elements.

For the Nluc-IRES-eGFP plasmid, the Nluc gene was amplified using the following primers: 5'-ACGACTCACTATAGGGAGACCCAAGCTTCGCCACCatggtcttcacactcgaagatttcg-3' and 5'-CGCCAGAATGCGTTCGC-3'.

The IRES fragment was amplified using the following primers: 5'-ggagtgaccggctggcggctgtgcgaacgcattctggcgtagacgttactggccgaagcc-3' and 5'-TTTTTCAAAGAAAACCACGTCCCC-3'.

The eGFP fragment was amplified using the following primers:
and 5'-acggggacgtggttttcctttgaaaaacacgataataccatggatggtgagcaagggcga-3' and 5'-GCGGCCGCCAGTGTGATGGATATCTGCAGAATTCctacttgtacagctcgtccatgccga-3'.

For the CD8LS-YeNL-CD4 plasmid, the YeNL gene was amplified from YeNL-pcDNA (Addgene plasmid no. 85201) using the following primers: 5'- gctgctccacgccgccaggccgGAATTCgtgagcaagggcgaggagct-3' and 5'-agacccgcctccgccCGCCA GAATGCGTTCGC-3'.

The CD4 fragment was amplified using the following primers:
5'-ggcggaggcgggtctttccagaaggcctccagcat-3' and 5'-GCCGCCAGTGTGATGGATATCTGCAGAATTCttattagcgccttcggtgcc-3'.

For the GeNL (Nluc-GF-mNeonGreen) pcDNA plasmid, the Nluc was amplified using the following primers:
5'-GGGCATGGACGAGCTGTACAAGggctttGAAGATTTCGTTGGGGACTGGC-3' and 5'-GCCGCCAGTGTGATG GATATCTGCAGAATTCctaCGCCAGAATGCGTTCGC-3'.

mNeonGreen was amplified using the following primers:
5'ATACGACTCACTATAGGGAGACCCAAGCTTCGCCACCatgGTGTCCAAGGGCGAAGAGGA-3' and 5'- CTTGTACAGCTCGTCCATGCCC-3'.

For the Fluc- and Akaluc-pET28a plasmids, the luciferase gene was amplified using the following primers:
5'-CGACTCACTATAGGGAGACCCAAGCTTATGGAAGATGCCAAAAACATTAAGAAG-3' and
5'-CACCGGCCTTATTCCAAGCGGCTTCGGCCAGTAACGTTTACACGGCGATCTTGCC-3'.

For the Nluc-pET28a plasmid, the Nluc gene was amplified using the following primers:
5'-GTTTAACTTTAAGAAGGAGATATACCATGGTAGTCTTCACACTCGAAGATTTCGTTG-3' and 5'-CTTTGTTAGCAGCCGGATTATTAGTGATGGTGATGGTGATGCGCCAGAATGCGTTCGCA-3'.

All PCR reactions (unless otherwise stated) were performed in a BioRad C3000 Thermocycler using the following conditions: 1) 95 °C for 3 min, 2) 95 °C for 30 s, 3) Tm of primers for 30 s, 4) 72 °C for 3 min, repeat steps 2-4 twenty times, then 72 °C for 5 min, and hold at 12 °C until retrieval from the thermocycler. Linearized vectors were generated via digestion with restriction enzymes *Hind*III and *Xho*I (New England BioLabs). The linearized vectors were combined with the appropriate insert by Gibson assembly (50 °C for 60 min). A portion of the reactions (3.0 µL) was directly transformed into TOP10 competent *E. coli* cells. Colonies containing the genes of interest were expanded overnight in 5 mL LB broth supplemented with ampicillin (100 µg/mL) or kanamycin (100 µg/mL) and DNA was extracted from colonies using a Zymo Research Plasmid Mini-prep Kit. Sequencing analyses were used to confirm successful plasmid generation.

**Microscopy**

Emission spectra for luciferase-expressing cells were recorded on a Zeiss LSM880 confocal microscope, equipped with a 32-channel spectral detector. Bioluminescent phasors were acquired on a Olympus IX83 TIRF microscope equipped with two Optosplit II (Cairn) image splitters and used in widefield mode. The light path is described in Supplementary Fig. S1. Briefly, collimated light exiting from the microscope body is split in half by a 50:50 beam splitter and sent to two Optosplit II image splitter devices. In each Optosplit II, the light is further split by a 50:50 beam splitter, half of which passes through a sine (or cosine) filter while the other half reaches the camera unfiltered. The signal ultimately reaches two identical sCMOS cameras (Prime 95b, Photometrics), the acquisition of which is controlled by the µManager software.

**In vitro bioluminescence imaging**

Phasor signatures for luciferase-luciferin pairs were recorded on the TIRF microscope using the 4-channel setup described above. For measurements with Fluc or Akaluc, D-luciferin, AkaLumine (AOBIOUS) or synthetic analogs (250 µM) were mixed with ATP (1 mM) in an Eppendorf tube and diluted to 200 µL with bioluminescence reaction buffer (20 mM Tris•HCl, 0.5 mg/mL BSA, 0.1 mM EDTA, 1 mM TCEP, 2 mM $MgSO_4$, pH = 7.8). For measurements with Nluc, YeNL or YNL, furimazine (Promega, 25–50 µM) or CTZ (Biosynth Carbosynth, 50 µM) was diluted to 200 µL with phosphate buffer (100 mM phosphate, pH 7.8). Purified luciferase (500 nM – 10 µM for Fluc, 10 nM for Nluc, YeNL or YNL) was added, and the solution was transferred to an 8-well chambered coverglass. Bioluminescence emission was recorded at room temperature using a 20x air objective (Zeiss UPlanSAPO 20x/0.75) with further 2x magnification. Acquisition times were set to 10 s per frame, and 20 frames total were collected per sample. Images were exported and analyzed as described below.

**In cellulo bioluminescence imaging**

HeLa cells were transfected with the appropriate reporter and plated on an 8-well chambered coverglass (Ibidi). Media was removed and replaced with a fresh stock of media (250 µL) containing furimazine (25–50 µM). Five minutes after the media change, the cells were imaged using a 20x air objective (Zeiss UPlanSAPO 20x/0.75) with further 2x magnification. For organelle-localized constructs, cells were imaged using a 60x oil objective (Zeiss Apo N 60x/1.49 Oil). All images were recorded with 10 s integration time and 20 frames total were collected per sample (unless otherwise stated). Images were exported and analyzed as described below. BRET efficiencies were calculated as the relative fraction of the acceptor ($f_2$) after unmixing the phasor, using the phasor of the donor and acceptor as pure components, as described below (Phasor unmixing section). The $f_2$ fraction corresponds to the fraction of photons emitted by the acceptor over the total number of photons emitted by both donor and acceptor.

**In vitro bioluminescence maxima**

Emission spectra for all luciferin analogs were recorded on an Agilent Cary Eclipse Fluorescence Spectrophotometer. Each luciferin (250 µM) was incubated in an Eppendorf tube with ATP (1 mM) and diluted to 1 mL with bioluminescence reaction buffer (20 mM Tris•HCl, 0.5 mg/mL BSA, 0.1 mM EDTA, 1 mM TCEP, 2 mM $MgSO_4$, pH = 7.8). Purified luciferase enzyme (10 µM) was added, and an aliquot (700 µL) was transferred to a 10 mm pathlength cuvette. The emission slit widths were set to 5. The detector gain was set to 500 mV. Emission data were collected at 1 nm intervals from 400−750 nm at ambient temperature. The acquisition times were set to 1 s/wavelength depending on the amount of light produced from each sample. Light emission was recorded as relative luminescence units (RLU), and the intensities were normalized to extrapolate emission maxima.

**Image processing**

Images were exported in TIFF format and processed with a custom MATLAB algorithm with the following workflow:

1. *Channel Splitting*: The images are split from a single TIFF file in 4 channels corresponding to the sine- and cosine-filtered channels and the appropriate reference channels. The images are cropped in regions of interest (ROIs) of approximately 900x385 pixels each and manually registered.

2. *Calibration*: For each day of experiment, a "Dark" image of only the camera noise is acquired as well as a "Bright" image of the transmission lamp (without sine-cosine filters) in order to characterize the camera offset and the effective light splitting in the four channels (described below, **Phasor transformation**). The files are cropped to the same ROIs as the experiments and the corrections are applied on a pixel-by-pixel basis.

3. *Processing*: Each intensity image is smoothed by a Gaussian filter with σ = 1 pixel and a variable number of frames (10 for organelles, 20 for cytoplasmatic reporters) is averaged together. Successively, a global user-defined threshold is set to remove the background contribution and the cells are segmented using the function "bwareafilt" in MATLAB and registered one at the time in all four channels using the function "imregister".

4. *Phasor transformation and clustering*: The four images are combined (see below, **Phasor transformation**) to compute the spectral phasor (g,s) coordinate on a pixel-by-pixel basis. Successively, the coordinates are clustered using a Gaussian-mixture model algorithm with the function "fitgmdist" and the clusters are assigned to the appropriate reporter based on the calibrated phasor position of the single cytoplasmatic reporters and color-coded accordingly.

5. *(Further processing)*: For organelle-targeting reporters, the images are blindly deconvolved with the function "deconvblind" using as initial guess a 5x5 matrix of ones. Successively, a median filter with a 41x41 pixels mask is applied and the resulting image is removed from the original image in order to remove the diffuse background contribution. The image is further smoothed with a Gaussian kernel with σ = 1 pixel and negative values set to 0.

**Phasor transformation and unmixing**

- *Dark calibration:* The file of the dark calibration is smoothed with a median filter with size 51x51 in order to remove high-frequency noise and an image $I_{i,dark}$ is stored for every channel. $i$ .

- *Bright calibration:* The file of the dark calibration is smoothed with a median filter with size 51x51 in order to remove high-frequency noise and an image $I_{i,bright}$ is stored for every channel $i$. In order to correct for the uneven intensity splitting in the sine (or cosine) channel and the reference intensity, two parameters are computed:

$$R_{COS} = \frac{I_{COS,bright} - I_{COS,dark}}{I_{INT(COS),bright} - I_{INT(COS),dark}}$$

$$R_{SIN} = \frac{I_{SIN,bright} - I_{SIN,dark}}{I_{INT(SIN),bright} - I_{INT(SIN),dark}}$$

Where $INT(SIN)$ and $INT(COS)$ refer to the reference channel for sine and cosine, respectively.

- *<u>Image correction:</u>* For the experiment files, each channel is corrected as follows

$$I_{INT(SIN)} = I_{INT(SIN)}^{raw} - I_{INT(SIN),dark}$$

$$I_{INT(COS)} = I_{INT(COS)}^{raw} - I_{INT(COS),dark}$$

$$I_{SIN} = \left(I_{SIN}^{raw} - I_{SIN,dark}\right) \cdot R_{SIN}$$

$$I_{COS} = \left(I_{COS}^{raw} - I_{COS,dark}\right) \cdot R_{COS}$$

Where the $raw$ denotes the uncorrected images.

- *<u>Phasor transformation:</u>* The (g,s) phasor coordinates are then obtained as follows:

$$g = 2 \cdot \left( \frac{\frac{I_{COS}}{I_{INT(COS)}} - F_{COS,min}}{F_{COS,max} - F_{COS,min}} \right) - 1$$

$$s = 2 \cdot \left( \frac{\frac{I_{SIN}}{I_{INT(SIN)}} - F_{SIN,min}}{F_{SIN,max} - F_{SIN,min}} \right) - 1$$

Where $F_{SIN,min}$ and $F_{SIN,max}$ correspond to the minimum and maximum transmission of the sine filter, as obtained from a transmittance measurement performed with a spectrophotometer. Similarly, $F_{COS,min}$ and $F_{COS,max}$ correspond to the minimum and maximum transmission of the cosine filter

- *<u>Phasor unmixing (2 components):</u>* given the phasor position of the two pure components $(g_1, s_1)$ and $(g_2, s_2)$, the phasor of a combination of the two will lie in position $(g, s)$ where $g = f_1 g_1 + f_2 g_2$ and $s = f_1 s_1 + f_2 s_2$. $f_1$ and $f_2$ are the relative contribution of the two pure components to the mixed signal, under the condition $f_1 + f_2 = 1$, that are retrieved by solving the linear system:

$$\begin{cases} g = f_1 g_1 + f_2 g_2 \\ s = f_1 s_1 + f_2 s_2 \end{cases}$$

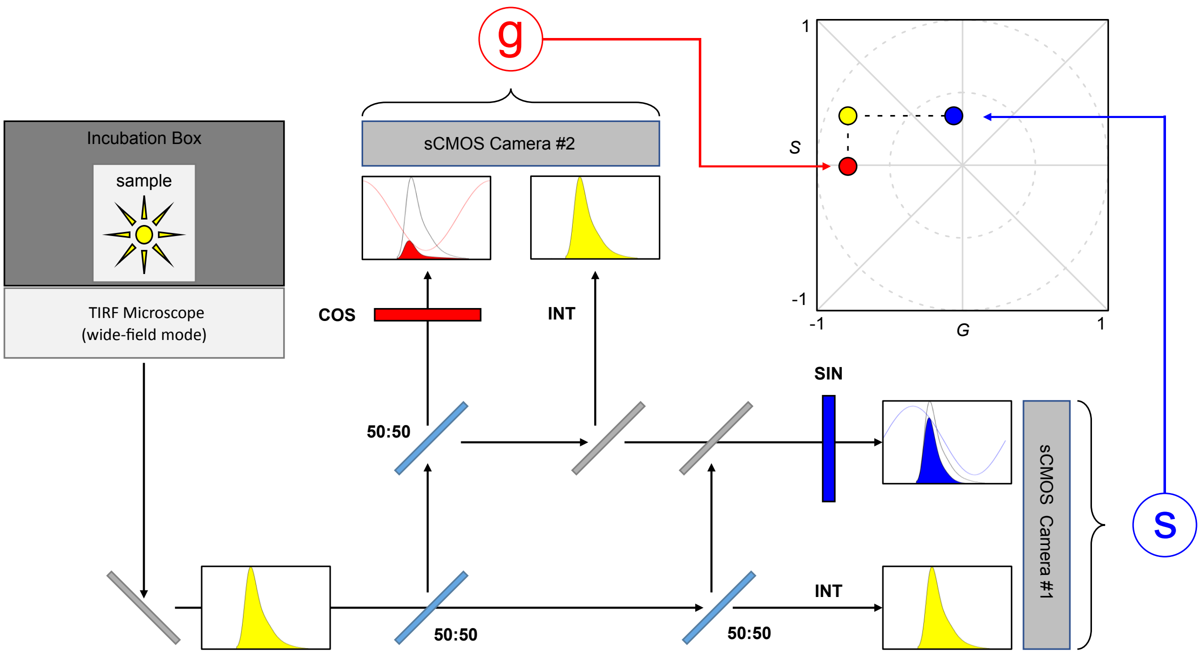


**Supplementary Fig. 1. Microscope setup for acquisition and optical transform of bioluminescent photons into phasor signatures.** Sample is placed in a temperature-, humidity- and $CO_2$-controlled box (Tokai) within the TIRF microscope, which is used in widefield mode. The luminescence emitted from the sample with a certain emission spectrum (yellow) is split in four identical channels via a series of 50:50 splitters (light blue) and mirrors (gray). The light is sent to two cameras that collect two channels each. Of the two channels, one is left unfiltered (yellow) while the other is filtered through either a sine filter (blue, camera #1) or a cosine filter (red, camera #2) and then integrated in the camera.[1] By referencing the sine and cosine images to the relative intensity images from the same camera, we can obtain (in a pixel-wise fashion) the g (cosine) and s (sine) coordinates that are successively used to identify the phasor transformation of the bioluminescent spectral emission (yellow dot, top right)[1-2]

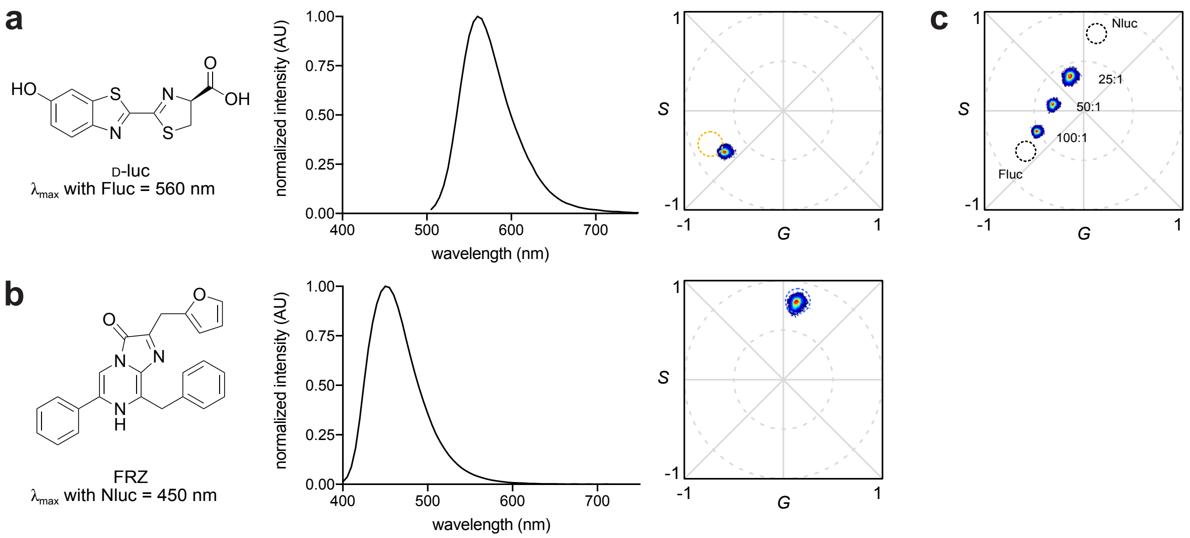


**Supplementary Fig. 2. Phasor signatures from distinct classes of bioluminescent probes.** Solutions of recombinant (a) firefly luciferase (Fluc, 500 nM) or (b) NanoLuc (Nluc, 10 nM) were mixed with D-luciferin (D-luc, 250 μM) and ATP (1 mM) or furimazine (FRZ, 50 μM),[3] respectively. Images were acquired using the microscope setup pictured in Supplementary Fig. 1 using a 10 s/frame integration time. A total of 20 frames were collected for each sample and the phasor locations were computed. The emission spectrum for each luciferase-luciferin pair is shown for reference (dashed circle). (c) Varying amounts of Fluc (500 nM) and Nluc (10 nM) were combined, and the mixtures were imaged with D-luc and FRZ. The molar ratios of Fluc:Nluc are shown. The relative contribution of Fluc in each sample was determined using the distance between the clusters on the phasor plot (36.5% in the 25:1 sample, 60.3% in the 50:1 sample and 82.7% in the 100:1 sample).

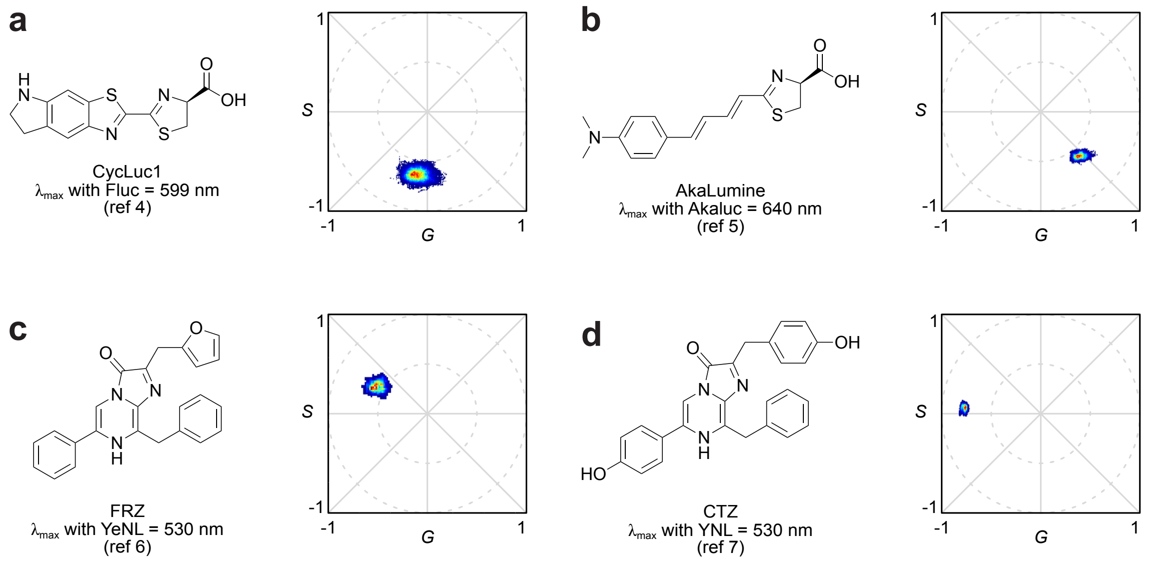


**Supplementary Fig. 3. Popular bioluminescent reporters produce distinct phasor signatures.** Solutions of recombinant (a) Fluc (500 nM), (b) Akaluc (10 μM), (c) yellow enhanced Nano-lantern (YeNL, 10 nM), and (d) yellow Nano-lantern (YNL, 100 nM) were mixed with their respective luciferin substrates (CycLuc1[4] or AkaLumine[5] = 250 μM with 1 mM ATP; FRZ[6] or CTZ[7] = 50 μM). Images were acquired using the microscope setup pictured in Supplementary Fig. 1 using a 10 s/frame integration time. A total of 20 frames were collected for each sample and the phasor locations were computed. The emission maximum for each luciferase-luciferin pair is shown for reference.

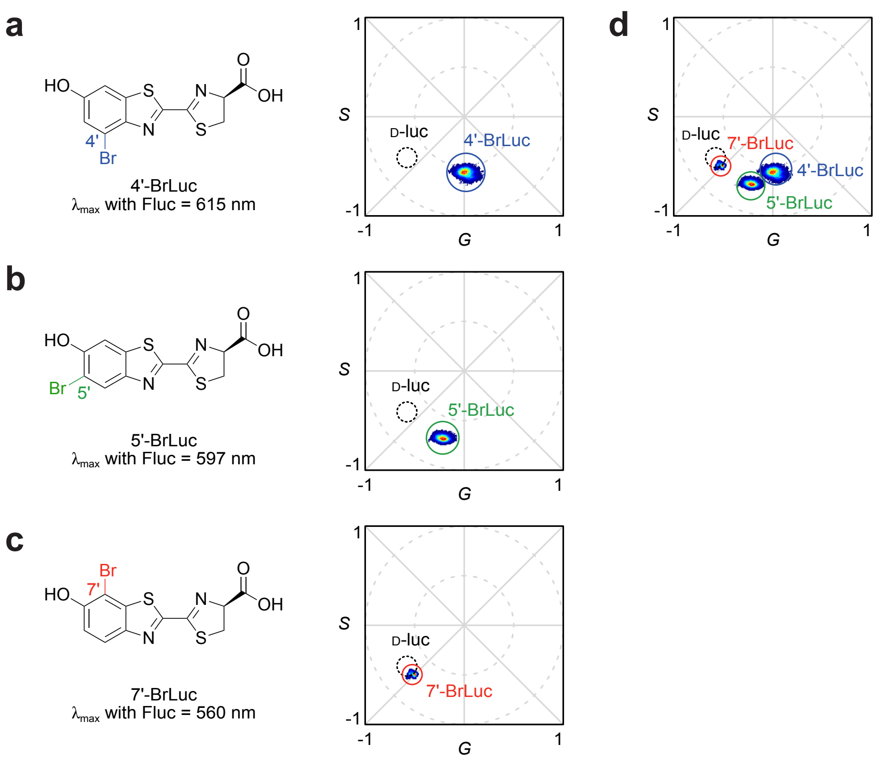


**Supplementary Fig. 4. Brominated luciferin isomers produce distinct phasor signatures.** A solution of recombinant Fluc (500 nM or 1 μM) was mixed with either (a) 4'-BrLuc, (b) 5′-BrLuc or (c) 7′-BrLuc (250 μM with 1 mM ATP).[8-9] Images were acquired using the microscope setup pictured in Supplementary Fig. 1 using a 10 s/frame integration time. A total of 20 frames were collected for each sample and the phasor locations were computed. The emission maxima for each substrate, and the phasor location for D-luciferin, are shown for reference. (d) Overlay of the phasor signatures from (a)-(c).

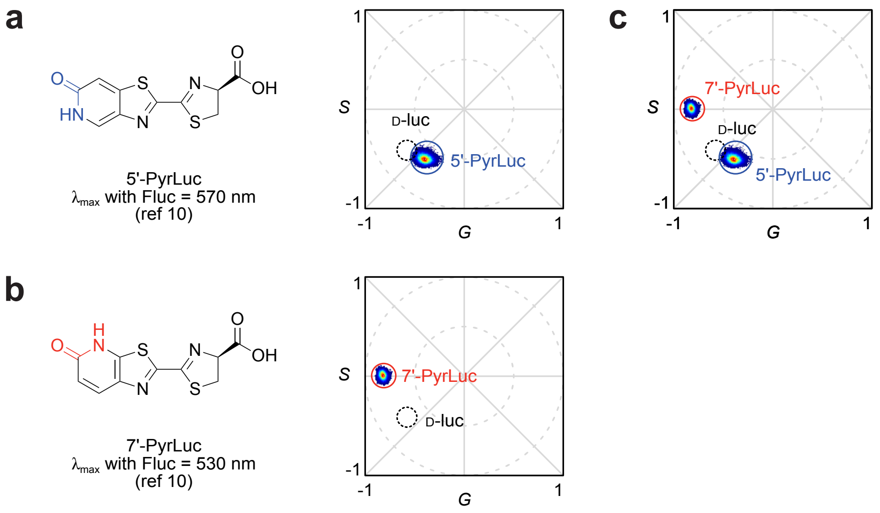


**Supplementary Fig. 5. Pyridone luciferin isomers produce distinct phasor signatures.** A solution of recombinant Fluc (1 or 10 μM) was mixed with either (a) 5′-PyrLuc or (b) 7′-PyrLuc (250 μM with 1 mM ATP).[10] Images were acquired using the microscope setup pictured in Supplementary Fig. 1 using a 10 s/frame integration time. The emission maxima for each substrate, and the phasor location for D-luciferin, are shown for reference. (c) Overlay of the phasor signatures from (a) and (b).

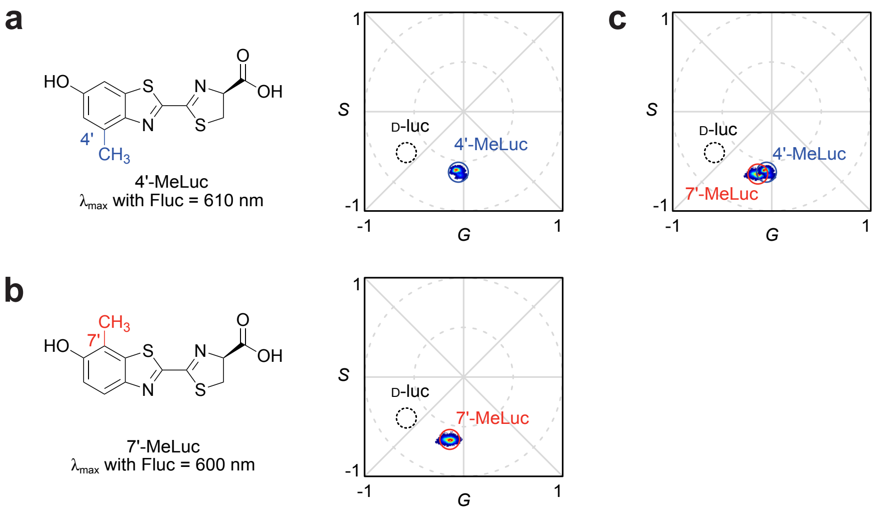


**Supplementary Fig. 6. Phasor signatures from Isomeric methylated luciferins.** A solution of recombinant Fluc (10 μM) was mixed with either (a) 4′-MeLuc or (b) 7′-MeLuc (250 μM with 1 mM ATP).[11] Images were acquired using the microscope setup pictured in Supplementary Fig. 1 using a 10 s/frame integration time. A total of 20 frames were collected for each sample and the phasor locations were computed. The emission maxima for each substrate, and the phasor location for D-luciferin, are shown for reference. (c) Overlay of the phasor signatures from (a) and (b).

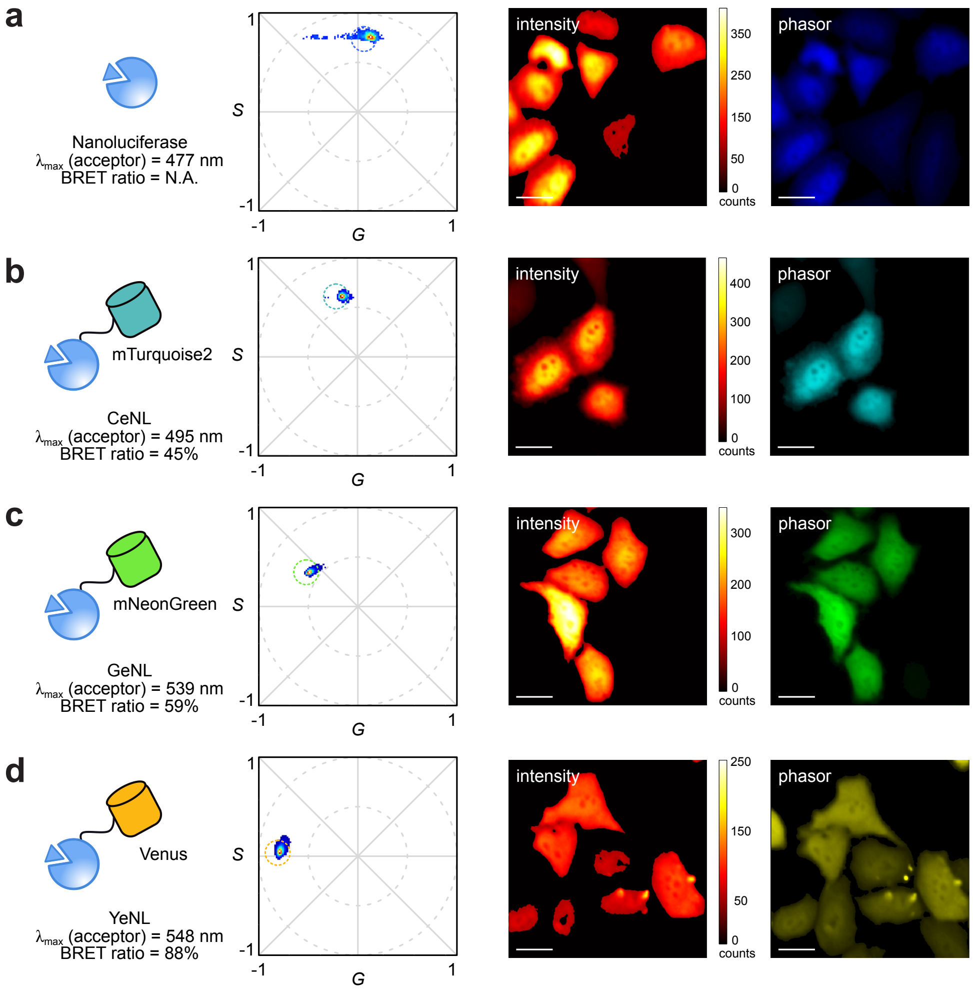


**Supplementary Fig. 7. Phasor signatures from BRET reporters.** HeLa cells were transiently transfected with plasmids encoding (a) Nluc, (b) CeNL, (c) GeNL, or (d) YeNL. The cells were then treated with FRZ (25–50 μM) ~36 h post transfection. Images were acquired using the microscope setup pictured in Supplementary Fig. 1 with a 20X air objective and 10 s/frame integration time. A total of 20 frames were collected for each sample and the phasor locations were computed. Each pixel in the resulting images was false colored according to the phasor signature. The expected phasor for each construct (based on its emission spectrum and BRET efficiency) is also shown for comparison. For all images, the scale bar = 25 μm. The images are color-coded according to the specific reporter, assigned after clustering.

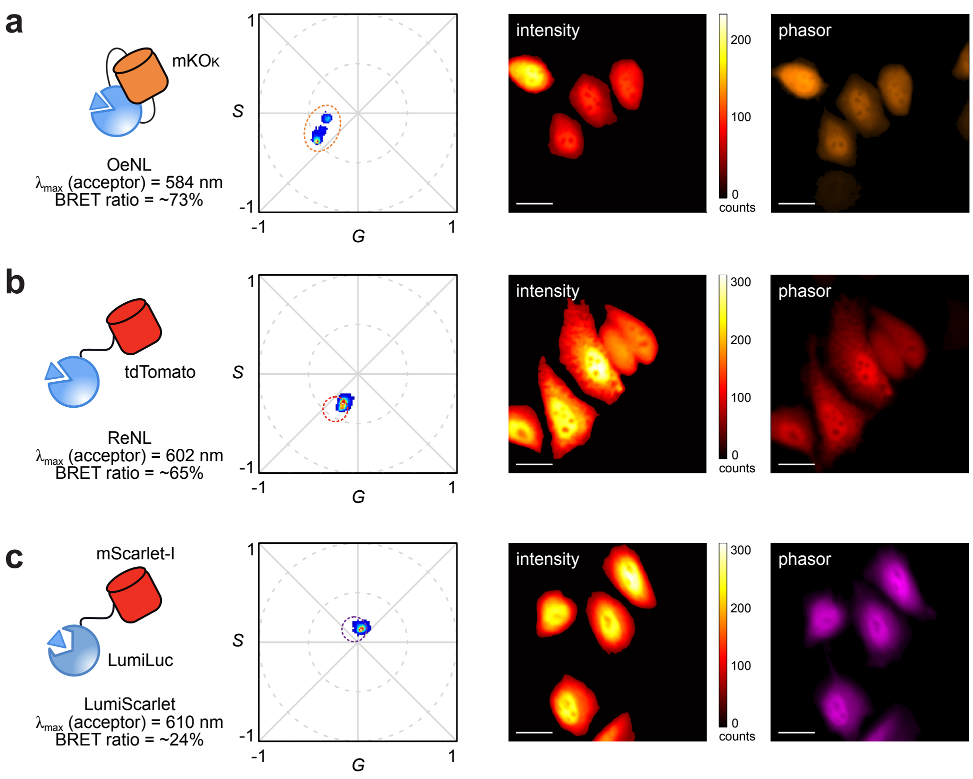


**Supplementary Fig. 8. BRET reporters with similar emission maxima can be distinguished via phasor analysis.** HeLa cells were transiently transfected with plasmids encoding (a) OeNL (b) ReNL, or (c) LumiScarlet. The cells were then treated with FRZ (25–50 μM) ~36 h post transfection. Images were acquired using the microscope setup pictured in Supplementary Fig. 1 with a 20X air objective and 10 s/frame integration time. A total of 20 frames were collected for each sample and the phasor locations were computed. Each pixel in the resulting images was false colored according to the phasor signature. The expected phasor for each construct (based on its emission spectrum and BRET efficiency) is also shown for comparison. For all images, the scale bar = 25 μm. The images are color-coded according to the specific reporter, assigned after clustering.

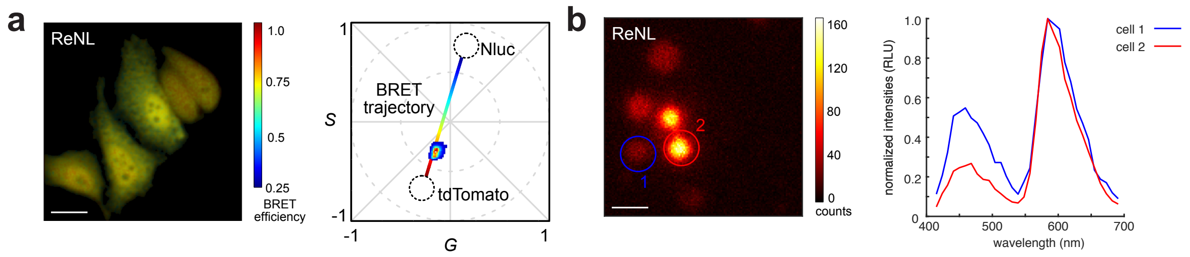


**Supplementary Fig. 9. Phasor signature of ReNL reveals cell-to-cell differences in BRET efficiency.** (a) Pixels from images acquired with ReNL-expressing HeLa cells (Supplementary Fig. 8b) were re-assigned based on the computed BRET efficiency. The donor location was assigned based on the phasor signature from Nluc-expressing cells (Supplementary Fig 7a), and the acceptor location was assigned based on the fluorescent spectrum of tdTomato. (b) BRET heterogeneity was validated on a confocal microscope (Zeiss LSM 880) equipped with a 32-channel spectral detector. For all images, the scale bar = 25 μm. The images are color-coded according to the specific reporter, assigned after clustering.

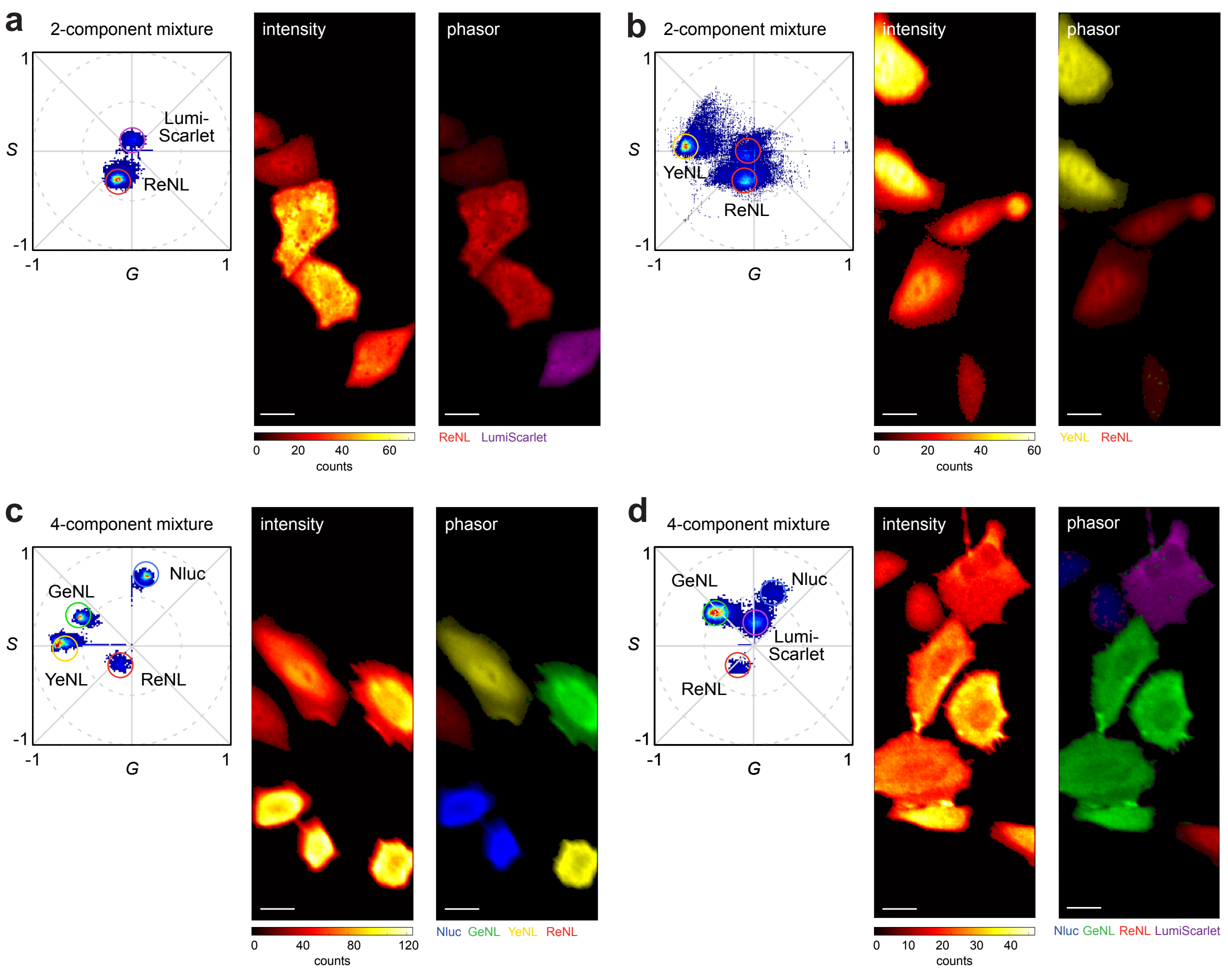


**Supplementary Fig. 10. Multicomponent imaging with BRET reporters.** HeLa cells were transfected with either Nluc or a BRET reporter (CeNL, GeNL, YeNL, ReNL, OeNL, or Lumi-Scarlet). The transfected cells were then randomly mixed and plated onto a glass slide 16 h post-transfection. The cells were treated with FRZ (25–50 μM). Images were acquired using the microscope setup pictured in Supplementary Fig. 1 with a 20X air objective and 10 s/frame integration time. A total of 20 frames were collected for each sample and the phasor locations were computed. Each pixel in the resulting image was false colored according to the phasor signature. Selected examples of (a, b) two-component, (c) four-component and (d) six-component mixtures are shown. Each component in the mixture was assigned by referencing to the single populations shown in Supplementary Figs. 7 and 8. For all images, the scale bar = 25 μm.

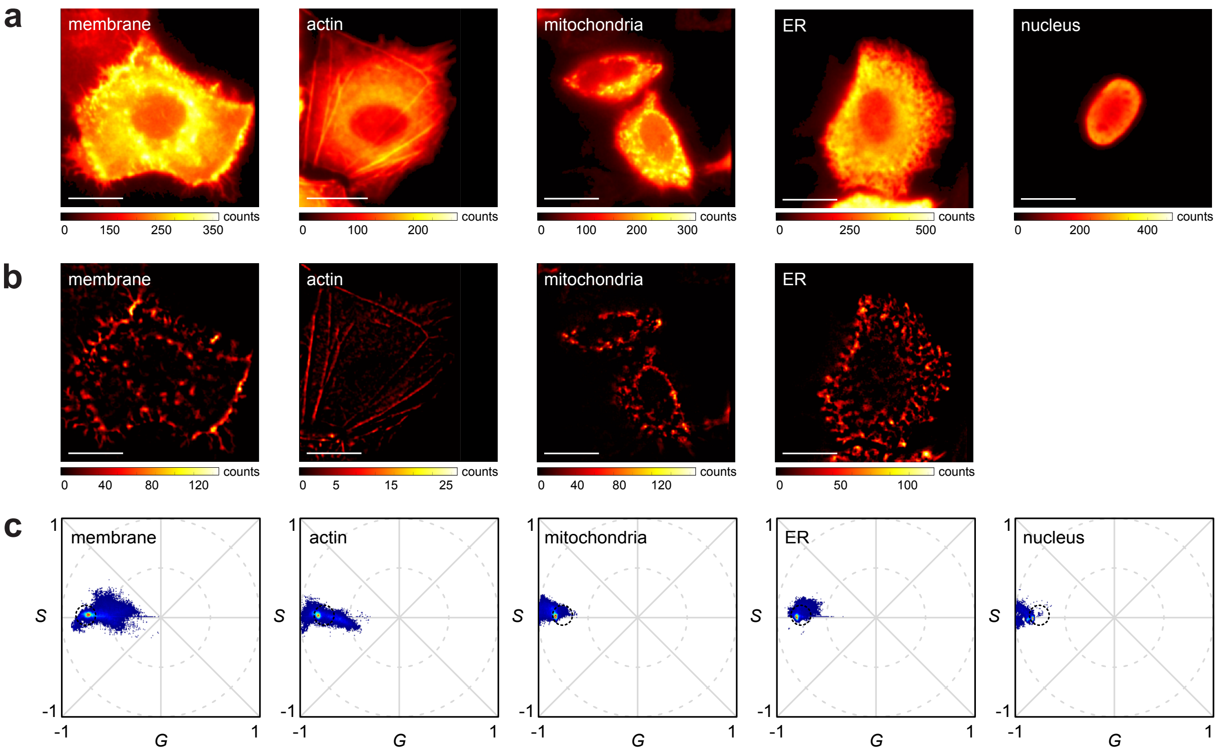


**Supplementary Fig. 11. Subcellular bioluminescent phasor imaging.** HeLa cells were transiently transfected with plasmids encoding YeNL localized to (from left to right): the membrane (CD8LS-YeNL-CD4), actin (YeNL-actin), mitochondria (Cox8x2-YeNL), ER (Carl-YeNL-KDEL) or nucleus (YeNL-H2B). The cells were treated with FRZ (25–50 μM) ~36 h post transfection. Images were acquired using the microscope setup pictured in Supplementary Fig. 1 with a 60x oil objective and 10 s/frame integration time. A total of 20 frames were collected for each sample and the raw intensity images (a) were subjected to image processing to remove out-of-focus signal. The resulting structural-enhanced images are shown in (b) and phasors computed for each construct are shown in (c). The phasor location of cytosolic YeNL is shown in black for comparison. For all images, the scale bar = 25 μm.

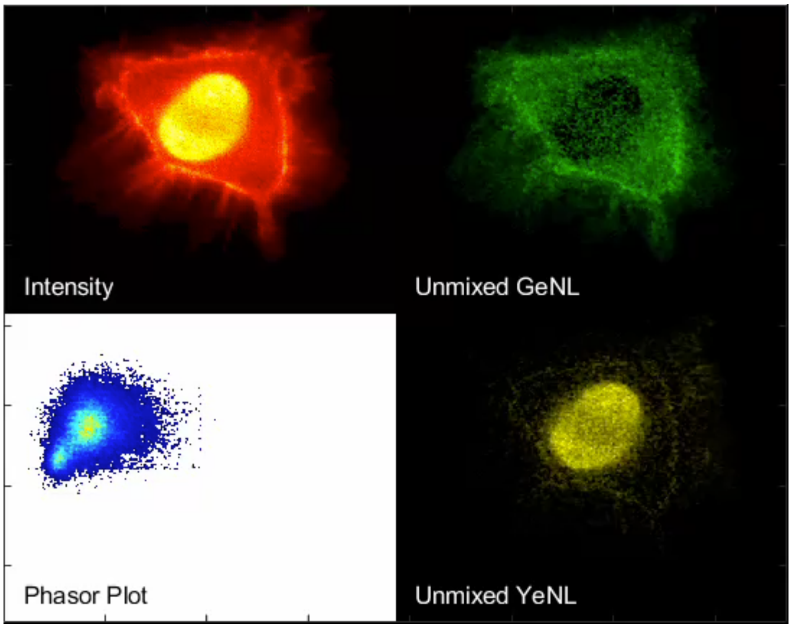


**Supplementary Video 1. Longitudinal multi-organelle tracking via bioluminescent phasor.** HeLa cells were transiently transfected with constructs comprising nuclear-localized YeNL and membraned-localized GeNL. The cells were treated with FRZ (25–50 μM) ~36 h post transfection. Images were acquired using the microscope setup pictured in Supplementary Fig. 1 with a 60x oil objective and 10 s/frame integration time. The sample was continuously imaged for 1 h (360 frames total). Selected snapshots at 0, 15, 30, and 45 min are shown in Fig. 2.
